# JMJD6 participates in the maintenance of ribosomal DNA integrity in response to DNA damage

**DOI:** 10.1101/833129

**Authors:** Jérémie Fages, Catherine Chailleux, Jonathan Humbert, Suk-Min Jang, Jérémy Loehr, Jean-Philippe Lambert, Jacques Côté, Didier Trouche, Yvan Canitrot

## Abstract

Ribosomal DNA (rDNA) is the most transcribed genomic region and contains hundreds of tandem repeats. Maintaining these rDNA repeats as well as the level of rDNA transcription is essential for cellular homeostasis. DNA damages generated in rDNA need to be efficiently and accurately repaired as rDNA repeats instability has been reported in cancer, aging and neurological diseases. Here, we describe that the histone demethylase JMJD6 is rapidly recruited at nucleolar DNA damage and is crucial for the relocalisation of rDNA in nucleolar caps. Yet, JMJD6 is dispensable for rDNA transcription inhibition. Mass spectrometry study revealed that JMJD6 interacts with the nucleolar protein Treacle and modulates its interaction with NBS1. Moreover, cells deficient for JMJD6 show increased sensitivity to nucleolar DNA damage as well as loss and rearrangements of rDNA repeats upon irradiation. Altogether our data reveals that rDNA transcription inhibition is uncoupled from rDNA relocalisation into nucleolar caps and that JMJD6 is required for rDNA stability upon the rDNA damage response through its role in nucleolar caps formation.

**Author summary:** Ribosomal DNA is the most transcribed genomic region composed of repeated sequences. Transcribed rDNA is essential for cellular homeostasis and cell proliferation. Numerous pathologies such as cancer and neurological disorders are described to present defective rDNA repeats maintenance. The mechanisms involved in the control of rDNA integrity involve major DNA repair pathways such as NonHomologous End-Joining and Homologous Recombination. However, how they are controlled and orchestrated is poorly understood. Here, we identified JMJD6 as a new member of the maintenance of rDNA integrity. We observed that JMJD6 controls the recruitment of NBS1 in the nucleolus in order to lead to the proper response to DNA damage at rDNA repeats.

**Author contributions:** YC and DT wrote the manuscript with input from their coauthors. JF, CC, JH, S-KM, JL, J-PL and YC performed the experiments. JC supervised JF and JH during purification and mass spectrometry analysis. YC and DT conceived the project, designed research and coordinated the studies.

## Introduction

Cells are continuously exposed to DNA damage which is repaired by different DNA repair pathways according to their specificities (double- or single-strand breaks, base modifications, …). DNA double-strand breaks (DSBs) are considered among the most deleterious DNA damage events and cells have evolved two main pathways to repair such damage: homologous recombination (HR) and non-homologous end joining (NHEJ)[1]. Moreover, the deleterious effect of DNA damage also depends on where the damage occurs within the genome with damage in transcribed regions being potentially the most detrimental. In this study, we focused on ribosomal DNA (rDNA) the most transcribed genomic locus. It is composed of tandem repeats (200-300) present on the five acrocentric chromosomes in human cells [2]. In interphase cells, DNA localises within nucleoli-specialized subcompartments in which rDNA transcription and rRNA processing take place.

Maintaining the number of rDNA repeats as well as the level of transcription is crucial for cellular homeostasis [3]. Instability of the rDNA repeats was reported in cancer [4] and Bloom and ataxia-telangectasia cell lines [5,6]. Therefore, DSBs generated in rDNA need to be efficiently and accurately repaired in particular to prevent recombination events that could result in loss of repeats in such repetitive regions. To date, the rDNA repair process is poorly understood and the mechanisms involved remain controversial [7,8]. Indeed, some studies have proposed that rDNA DSB repair could be mediated by NHEJ within the nucleolus [9,10]. However, persistent breaks have been shown to be relocalised to nucleolar caps at the periphery of the nucleolus. This process is ATM-dependent and has been proposed to be tightly linked to transcription inhibition [9,10,11,12]. Those breaks directed to nucleolar caps may possibly be more accessible to nucleoplasmic repair proteins and may be repaired by the action of HR proteins shown to be present at the nucleolar periphery throughout the cell cycle [13]. Lastly, a distinct nucleolar DNA damage response involving an ATM-TCOF1-MRN axis was identified to mediate rDNA transcription inhibition concomitantly to nucleolar restructuring in response to rDNA induced DSB [12].

DNA repair is influenced by the chromatin context marked by numerous histone post translational modifications [14,15,16]. In this study, we identify JMJD6, a member of the histone demethylase family, as an important player of the response to DNA damage occurring in rDNA. JMJD6 is a member of the Jumonji C domain-containing protein family and has been described as presenting dual enzymatic activity, an arginine demethylase contributing to histone methylation control [17] and a lysyl hydroxylase activity targeting non histone proteins such as p53 [18] and the splicing factor U2AF65 [19]. Recently, an additional function for JMJD6 as a tyrosine kinase targeting the Y39 tyrosine of the histone variant H2A.X has been reported in triple negative breast cancer cell lines overexpressing JMJD6 [20]. The recent study by Liu et al [20] describes JMJD6 as responsible for such a phosphorylation and raises the question of the role of JMJD6 in the DNA damage response. More recently, JMJD6 was shown to modulate the DNA damage response independently of its enzymatic activity through the modulation of histone H4 acetylation with its depletion leading to increased DNA double strand breaks repair and resistance to ionizing radiations [21]. Here we show that JMJD6 is recruited to DNA damage generated in the nucleolus and influences DNA repair, increasing genomic instability within rDNA when JMJD6 is defective. Furthermore we show that JMJD6 is necessary for the relocalisation of the repair factor NBS1 in the nucleolus upon DNA damage, thus providing novel insight into the mechanism underlying repair of rDNA DSBs.

## Results

### JMJD6 is involved in the DNA damage response to ionizing radiation

In order to obtain an integrated view of the involvement of histone demethylases in the response to DNA damage, we performed a screen using a siRNA library directed against all known or putative (Jumonji domain-containing proteins) histone demethylases in U2OS cells. Using as a read-out the DNA damage γH2AX foci formation, we identified JMJD6 as a hit whose depletion caused higher γH2AX staining following ionizing radiation (IR) exposure. To confirm these observations and to rule out possible off target effects, we used two additional siRNAs directed against JMJD6, which efficiently decreased JMJD6 expression (Figure 1A). We observed that transfection of any of these siRNAs increased γH2AX staining both before and in response to IR (Figure 1B and C). Taken together, this data indicates that JMJD6 is required for the normal response to irradiation-induced DNA damage in U2OS cells. Using live cell imaging, we next tested whether the effect of JMJD6 on DNA damage signaling and repair was direct by assessing the recruitment of GFP-tagged JMJD6 to sites of laser induced-DNA damage. We consistently observed a strong enrichment of JMJD6-GFP at the site of DNA damage in the nucleolus (Figure 1D) which was not detected when using 53BP1-GFP, as a positive control, or GFP alone as a negative control (Figure S1). In addition, the recruitment of JMJD6-GFP at DNA breaks in the nucleolus was quicker (around 1-2min) than that of 53BP1 in the nucleoplasm (10 min). Of note, in some cells expressing GFP-tagged JMJD6, we found some enrichment of JMJD6 at nucleoplasm sites of DNA damage (Figure S1).

**Figure 1.**
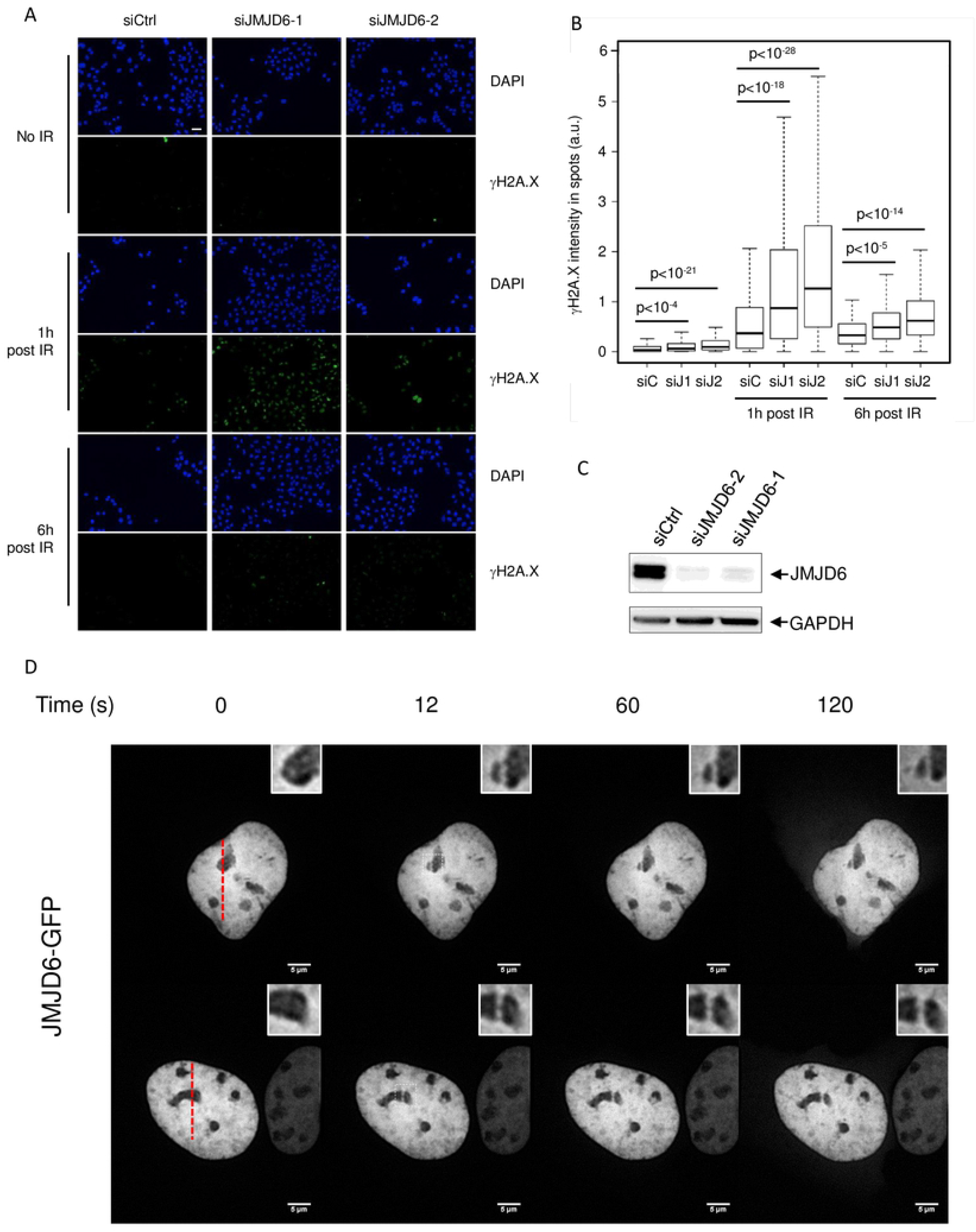
JMJD6 expression is required for the normal DNA damage response to ionizing radiation exposure. A. siRNA efficacy tested through Western blot analysis on whole cell extracts. B. Images of U2OS cells transfected with the indicated siRNA, exposed to ionizing radiations (8 Gy) and subjected to DAPI and gH2AX staining one hour or six hours following irradiation, as indicated. Scale bar 50 µm. C. Quantification of gH2AX staining by high throughput microscopy in cells from A. A minimum of 200 cells were quantified for each conditions. A representative experiment from 3 is shown. The p values of the difference between the indicated samples are shown (Wilcoxon test). D. U2OS cells transfected by a plasmid expressing JMJD6-GFP fusion protein and subjected to local laser irradiation 24h later. Images of cells before and after the indicated time following nucleus laser irradiation are shown. The red dashed lines represent the laser irradiated region in the nucleus. insert: magnification of the nucleolus irradiated regions. Scale bar 5 µm.

Note that this was not observed in all cells which contrasts with the fact that JMJD6-GFP was systematically recruited to nucleoli upon damage when damage are inflicted on nucleolus. Thus, this data indicates that JMJD6 is rapidly recruited to DNA damage sites which occur in the nucleolus, strongly suggesting that it could play a direct role in DNA damage signaling and repair following breaks occurring on nucleolar DNA. Recruitment of JMJD6 to DSB was also investigated by performing ChIP experiment. We constantly observed the presence of JMJD6 at rDNA in the absence of break induction. However, we did not observe any further enrichment in the vicinity of induced-DSB (Figure S2). This result could be due to a rapid and transient recruitment of JMJD6 to DSBs at the rDNA. Because of the repetitive nature of rDNA, the JMJD6 presence to uncut repeats could mask the recruitment at few copies induced following breaks. Nevertheless, it shows a recruitment of JMJD6 at rDNA, consistent with a role in rDNA damage management.

### Cells deficient for JMJD6 are sensitive to DNA damage occurring within rDNA

We next tested whether JMJD6-depleted cells, which present an impaired DNA damage response, were sensitive to irradiation by performing clonogenic cell survival assays post-irradiation. We generated JMJD6 KO cell lines (JMJD6 -/-) that were found to be more sensitive to IR than parental cells (Figure 2A and Figure S3). Importantly, JMJD6 KO cells complemented with the expression of JMJD6 from a plasmid were less sensitive than KO cells, regaining nearly equivalent sensitivity to IR than parental U2OS cells (Figure 2A and Figure S3). Thus, this data indicates that the defect in the DNA damage response observed upon JMJD6 knock down using siRNA translates into increased sensitivity to DNA damage in JMJD6-negative cells. In addition, cells complemented with a catalytic inactive form of JMJD6 (KO+Mut) showed no complementation for the sensitivity to IR (Figure 2A) although it was recruited at laser-induced damage as observed for the WT form of JMJD6 (Figure S4). The recruitment of JMJD6 at nucleolar damage sites indicates its potential involvement in rDNA damage response and repair. Considering the size of the rDNA loci, the irradiation dose we used in the clonogenic cell survival assay should not lead to breaks in the rDNA in a significant proportion of cells. Therefore, to test whether JMJD6 depletion could lead to increased sensitivity to damage generated specifically in the nucleolus, we carried out a cell survival assay following targeted induction of rDNA breaks using CRISPR-cas9 and a guide RNA targeting rDNA [13]. In this experiment, we included a control guide RNA targeting the ATP1A1 gene coding for the Na/K pump together with a donor DNA, which renders genome-edited cells resistant to ouabain [22] (Figure 2B). By performing clonogenic experiments in the presence of ouabain, we could thus rule out any effect of JMJD6 depletion on Cas9 or guide RNA expression. In the absence of control guide RNA targeting the ATP1A1, ouabain treatment killed all cells (data not shown), whereas transfection of this guide led to the generation of many ouabain resistant clones, reflecting targeted genome editing at the ATP1A1 gene (Figure 2C, left). Co-transfecting the guide RNA targeting rDNA strongly diminished the number of ouabain-resistant colonies (Figure 2C, right), indicating that DSBs in rDNA impaired cell survival. Interestingly, depletion of JMJD6 further decreased cell survival upon CRISPR-induced rDNA breaks (Figure 2C and D). Taken together, this data indicates that the full expression of JMJD6 is specifically required for the management of DSBs occurring at the rDNA locus, consistent with our finding that JMJD6 is always recruited to the nucleolus upon DNA damage.

**Figure 2.**
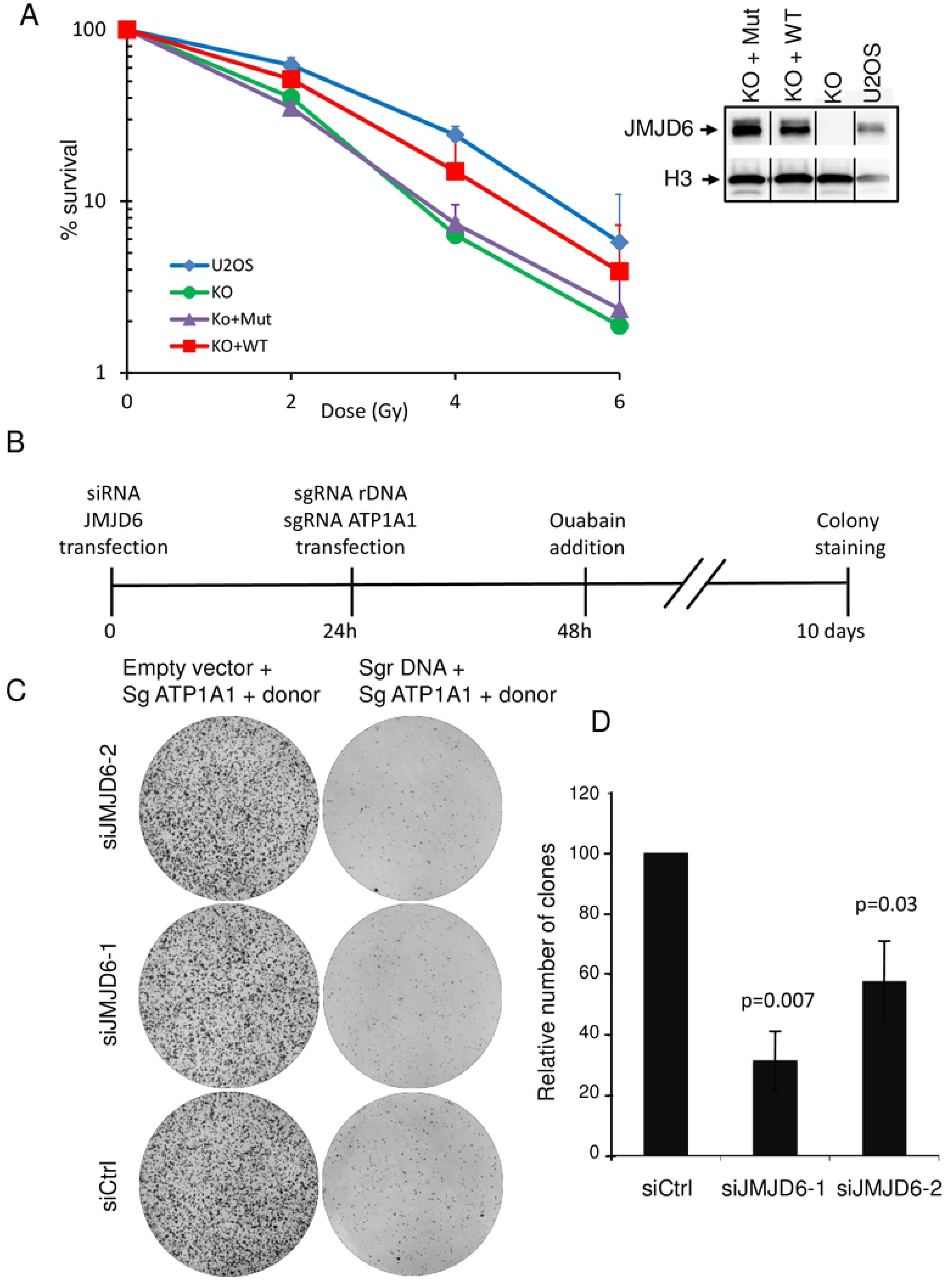
JMJD6-depleted cells are more sensitive to DSBs induced in rDNA. A. Clonogenic cell survival assay performed on U2OS cells, JMJD6-KO and JMJD6-KO-complemented cell lines with WT (KO+WT) and catalytic inactive forms of JMJD6 (KO+Mut) and exposed to increased doses of irradiation. Expression of JMJD6 in the different cell lines examined by Western blot. The bar indicates that the original image was cut. B. Scheme of the CRISPR experiment designed to test the sensitivity of JMJD6 depleted cells subjected to targeted rDNA breaks. U2OS cells were transfected with a vector expressing a guide RNA targeting the rDNA (sgrDNA) together with a guide RNA targeting the ATP1A1 gene (sgRNA ATP1A1) and a donor DNA rendering ATP1A1 resistant to ouabain. As a control an empty vector not coding for guide RNA targeting rDNA but with a guide RNA targeting the ATP1A1 gene (sgRNA ATP1A1) and a donor DNA rendering ATP1A1 resistant to ouabain was used. 10 days later, clones were stained with crystal violet. C. Example of a typical experiment. D Quantification of the experiment, the number of clones was calculated relative to 100, with 100 corresponding to cells transfected by the control siRNA. The mean and standard deviation from three independent experiments are plotted. The p values of the difference between the siJMJD6 samples and the Ctrl siRNA sample are indicated (Student t test).

### JMJD6 depletion generates genetic instability at the rDNA level

Faithful repair of DSBs occurring in rDNA allows the maintenance of rDNA repeats integrity. rDNA repeats are present on the five acrocentric human chromosomes and can be visualized in metaphase as 10 UBF foci, which correspond to the Nucleolus Organizer Regions (NORs) (Figure 3A) [9,13]. To study the involvement of JMJD6 to preserve rDNA stability JMJD6 KO or control cells were subjected or not to IR (2Gy) followed by a 30h recovery period before scoring the number of NORs using UBF as a marker (Figure 3A). As expected, the median number of NORs in control cells was around 10. In JMJD6 KO cells, a significant decrease was already observed in absence of DNA damage (median number of 9) meaning that JMJD6 is necessary even in absence of external DNA damage. After IR exposure an additional loss of NORs was observed (Figure 3B). This decrease in NOR number corresponds to *a bona fide* loss of rDNA sequences since UBF expression was unchanged (Figure 3C). Moreover, it does not result in an off-target effect of the KO cell line since the number of NORs was largely preserved in JMJD6 complemented-KO cells (Figure 3B). Thus, this data indicates that JMJD6 expression is required for the maintenance of the number of NORs upon irradiation, indicating its major role in the maintenance of rDNA repeats integrity. To better characterized the genetic instability at rDNA in response to DNA damage we performed DNA FISH combing using fluorescent FISH probes targeting rDNA (Figure 3D). Example of normal rDNA repeats organized in head-to-tail (regular succession of red-green units) is shown and ascribed as canonical. Example of rDNA rearrangements identified as non-canonical are pointed by a star. Results show that in absence of external DNA damage JMJD6 depletion *per se* induced a higher level of rDNA rearrangements (Figure 3E and Figure S5). In response to induced-DSB we observed an increase in rDNA rearrangements in control cells which was higher in JMJD6-depleted cells. Together these results confirm that JMJD6 is important to preserve rDNA from major rearrangement.

**Figure 3.**
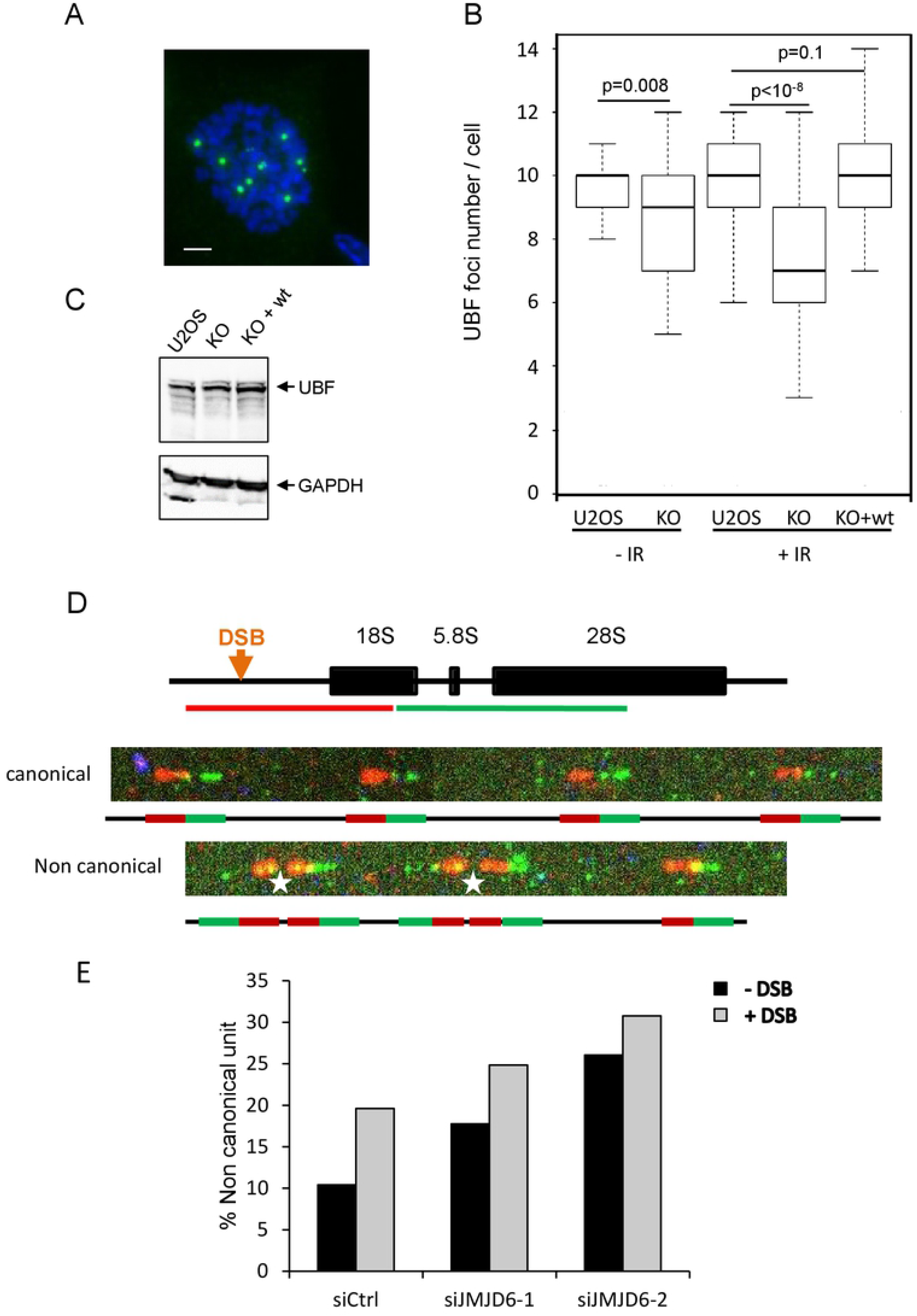
JMJD6 expression is required for rDNA repeat integrity following DNA damage. A. Representative image showing individual NORs in a U2OS cell in metaphase stained using an anti UBF antibody. Scale bar 5 µm. B. Ionizing radiation (2 Gy) exposure of U2OS cells, U2OS cells inactivated for JMJD6 expression (KO) and a clone from the latter cell line in which wild type JMJD6 was reintroduced (KO + wt). The number of UBF foci in cells was then counted and the results represented as box plot. For each point a minimum of 50 metaphases were scored. Results from one representative experiment from 2 independent experiments is shown. The p values of the difference between the indicated samples are shown (Wilcoxon test). C. Western blot analysis of UBF expression in the different cell lines. D. Evaluation of rDNA rearrangements by FISH combing. Representation of a rDNA repeat with the position of the DSB induced after OHtam treatment. The green and red lines represent the FISH probes used in the DNA FISH combing experiment. Examples of canonical (without rDNA rearrangement) and non-canonical (with rDNA rearrangement indicated by a star) rDNA repeats. E. Quantification of non-canonical rearrangements measured before and after DSB induction in siRNA control and siRNA JMJD6-depleted cells. Quantification was performed on duplicate samples with more than 400 units examined on each samples. One representative experiment from two independent experiments is shown.

### JMJD6 controls the NBS1-Treacle interaction in response to DNA damage

We next investigated the mechanism by which JMJD6 could influence DNA repair in the nucleolus. To characterize the JMJD6 physiological interactome, we raised a K562-derived cell line in which the endogenous JMJD6 gene was edited with CRISPR/cas9 to produce a C-terminal 3xFlag/2xstreptavidin-tagged JMJD6 protein. After tandem affinity purification from nuclear soluble extracts, we examined JMJD6-interacting proteins by mass spectrometry. Consistent with the known role of JMJD6 in mRNA splicing [23,24], we recovered many proteins linked with the spliceosome apparatus and mRNA 3’-end processing (CPSF, SYMPK, DDX41, WDR33), validating our experimental strategy. In addition, our results showed proteins specific of the nucleolar compartment and associated with rDNA transcription (Figure 4A). Among them, TCOF1 (Treacle) drew our attention as it participates in the response to DNA damage in the nucleolus [25,26]. We confirmed the JMJD6-TCOF1 interaction in U2OS cells by performing Proximity Ligation assay (PLA) (Figure 4B-C) together with confocal microscopy showing co-localisation in the nucleolus (Figure S6).

**Figure 4.**
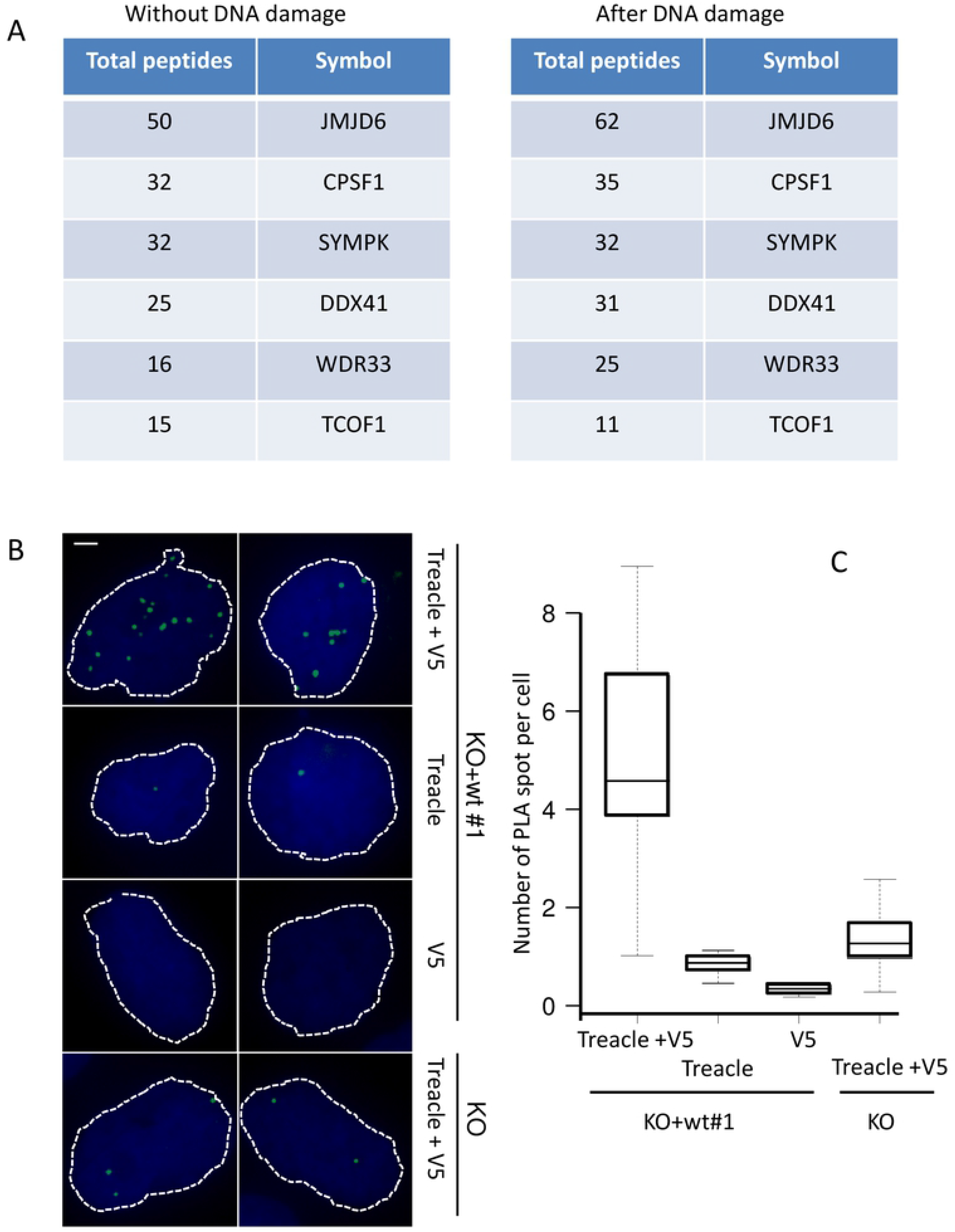
JMJD6 interacts with TCOF1 (Treacle). A. List of JMJD6 interactors identified after mass spectrometry (total spectral counts) after tandem affinity purification of endogenous JMJD6 isolated from K562 cells before and after DNA damage induced with etoposide (20µM 1h). B. Images of JMJD6-V5/Treacle (TCOF1) interaction revealed by Proximity Ligation Assay (PLA). Scale bar 5 µm. C. Quantification of PLA-induced spot for the different conditions of antibodies was performed on at least 200 cells for each conditions.

Interestingly, TCOF1 has been reported to be essential for the relocalisation of NBS1 into the nucleolus in response to DNA damage to repress rDNA transcription [25,26]. We thus performed a PLA assay for monitoring Treacle-NBS1 interaction after DNA damage according to JMJD6 status. Using the JMJD6-KO cell lines and the WT or Mutant -complemented cell lines, we observed a higher level of interaction between NBS1 and Treacle in JMJD6 KO and Mutant-complemented cell lines compared to WT-complemented cell line after IR exposure (Figure 5 and Figure S7). The proximity of the 3 proteins was confirmed by confocal microscopy in NBS1-GFP transfected cells (Figure S8). These results show that JMJD6 could control or influence the NBS1-Treacle interaction.

**Figure 5.**
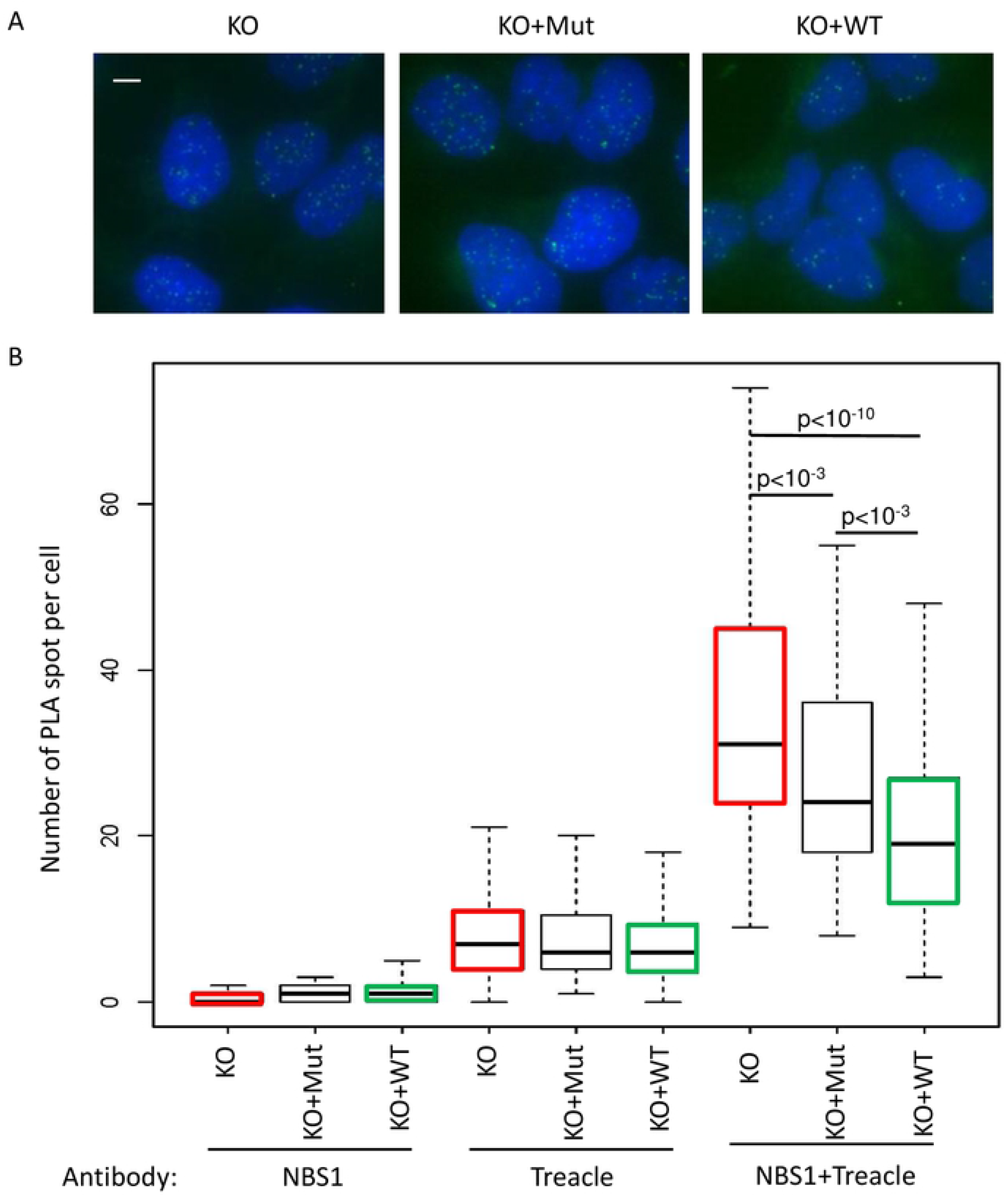
JMJD6 controls the interaction between Treacle and NBS1. A. Images of NBS1/Treacle interaction revealed by Proximity Ligation Assay (PLA) in cells exposed to IR. Scale bar 5 µm. B. Quantification of PLA-induced spot for the different conditions of antibodies was performed on at least 200 cells for each conditions. One representative experiment from two independent experiments is shown. The p value of the difference between KO and KO+Mut to KO+WT are shown.

### JMJD6 depletion does not affect the rDNA transcription response upon DNA DSBs induced in rDNA

The TCOF/NBS1 complex has been shown to mediate transcriptional silencing of rDNA repeats upon DNA breaks induced at rDNA. To analyse the effect of JMJD6 depletion on rDNA transcription we used the DIvA cell line in which DSBs are produced across the genome by the AsiSI endonuclease [27] one of which being located within rDNA and potentially generating one DSB per rDNA repeat [28]. As expected, control cells showed a decrease in rDNA transcription following DSB induction, as measured by the incorporation of 5-FUrd metabolic labelling (Figure 6A and B). Strikingly, rDNA transcription was further decreased in JMJD6-depleted cells compared to control cells (Figure 6) while rDNA transcription in absence of DNA damage remained largely unaffected (although in some experiments such as the one shown in Figure 6C, we could observe a slight decrease of basal rDNA transcription upon JMJD6 knock-down). Similar effects on rDNA transcription repression were observed 1h post IR (8 Gy), with a further reduction of transcription in JMJD6 depleted cells (Figure 6C and D). Interestingly, rDNA transcription levels had fully recovered 6 hours following IR in both control and JMJD6-depleted cells (Figure 6C and D), indicating that the rDNA transcription decrease observed upon JMJD6 depletion was transient. Altogether, this data shows that JMJD6 expression defect does not perturb the transcriptional repression of rDNA observed after DNA damage induction.

**Figure 6.**
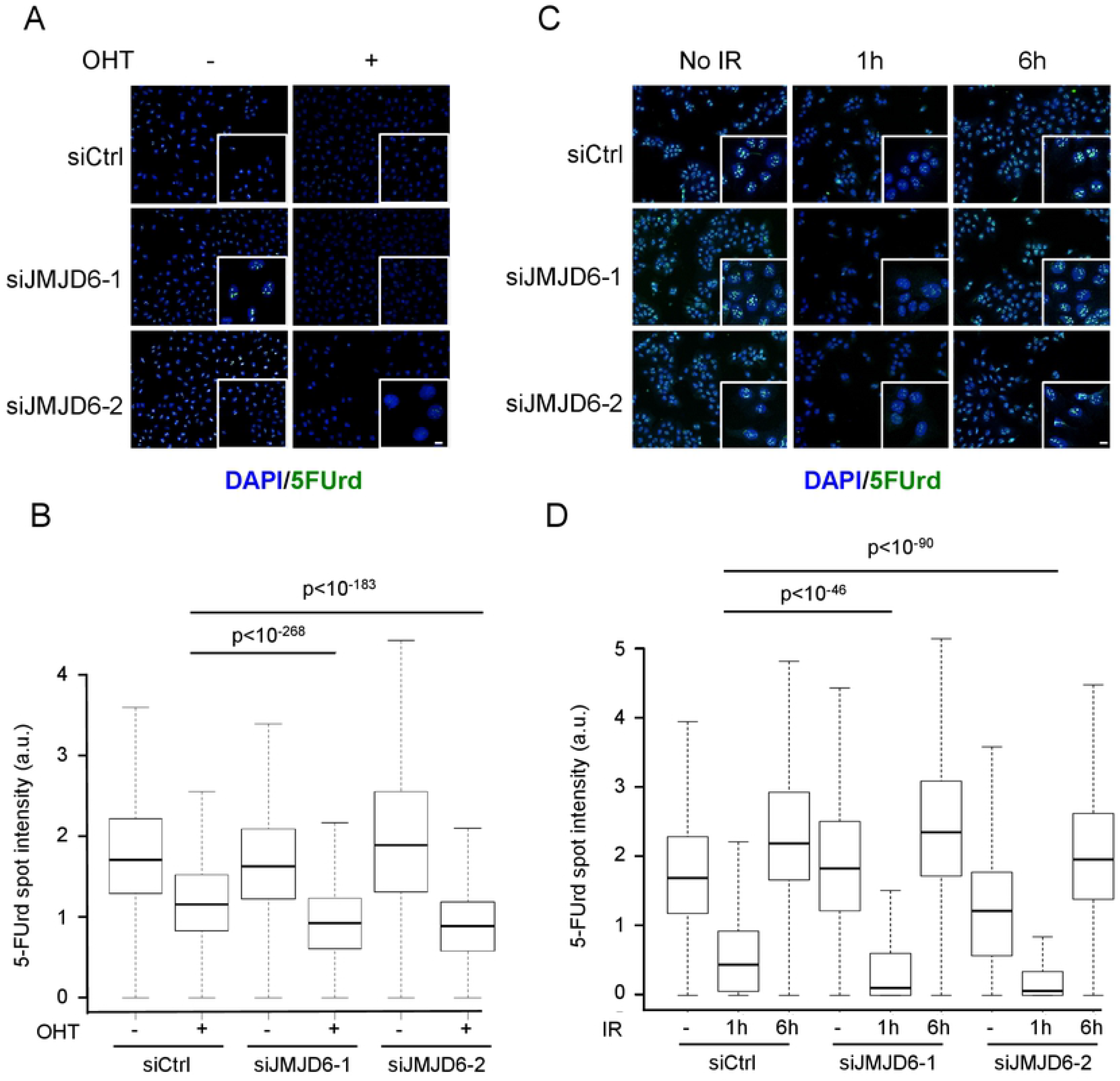
JMJD6 expression favours rDNA transcription following DNA damage. A. Level of rDNA transcription assessed through 5FUrd incorporation following endonuclease-mediated DSBs (+OHT) in U2OS DIVA cells transfected with the indicated siRNA, insert: a magnified region of the image. Scale bar 5 µm. B. Quantification of 5FUrd staining by high throughput microscopy in cells from A. A representative experiment from 2 independent experiments is shown. The p values of the difference between the indicated samples are shown (Wilcoxon test). C. Same as in A, except that cells were exposed to ionizing radiation (IR) (8 Gy) or not and stained 1 hour or 6 hours following irradiation. Scale bar 5 µm. D. Quantification of 5FUrd staining by high throughput microscopy in cells from C. A representative experiment from 2 independent experiments is shown. For each data point a minimum of 200 cells were quantified. The p values of the difference between the indicated samples are indicated (Wilcoxon test).

### JMJD6 is required for nucleolar caps formation

The Treacle/NBS1 complex was also shown to be important for the relocalisation of rDNA at the nucleolar periphery in structure called nucleolar caps, upon induction of DSBs in rDNA [9,13]. We thus tested the involvement of JMJD6 in this process by detecting nucleolar caps using UBF staining [28]. As expected, upon DSB induction in rDNA using DIvA cells, nucleolar caps were readily formed in a significant proportion of cells (Figure 7A). Strikingly, less cells displaying nucleolar caps were observed in JMJD6 depleted cells compared to control cells (Figure 7B). This decrease in nucleolar caps formation was not caused by an altered expression of UBF (Figure 7C), nor by changes in cell cycle distribution (Figure S9). Similar results were obtained after exposure to a high dose of IR (20 Gy 6h) showing that defective generation of nucleolar caps in JMJD6 depleted cells was not restricted to endonuclease-generated DNA damage (Figure S10). This data indicates that JMJD6 expression is required for the formation of nucleolar caps. Importantly, nucleolar caps have been proposed to be associated with rDNA transcription inhibition in response to DNA damage [11]. However, here we show that nucleolar caps formation and transcription inhibition can in fact be uncoupled since JMJD6 depleted cells display transcriptional repression, with yet less nucleolar caps.

**Figure 7.**
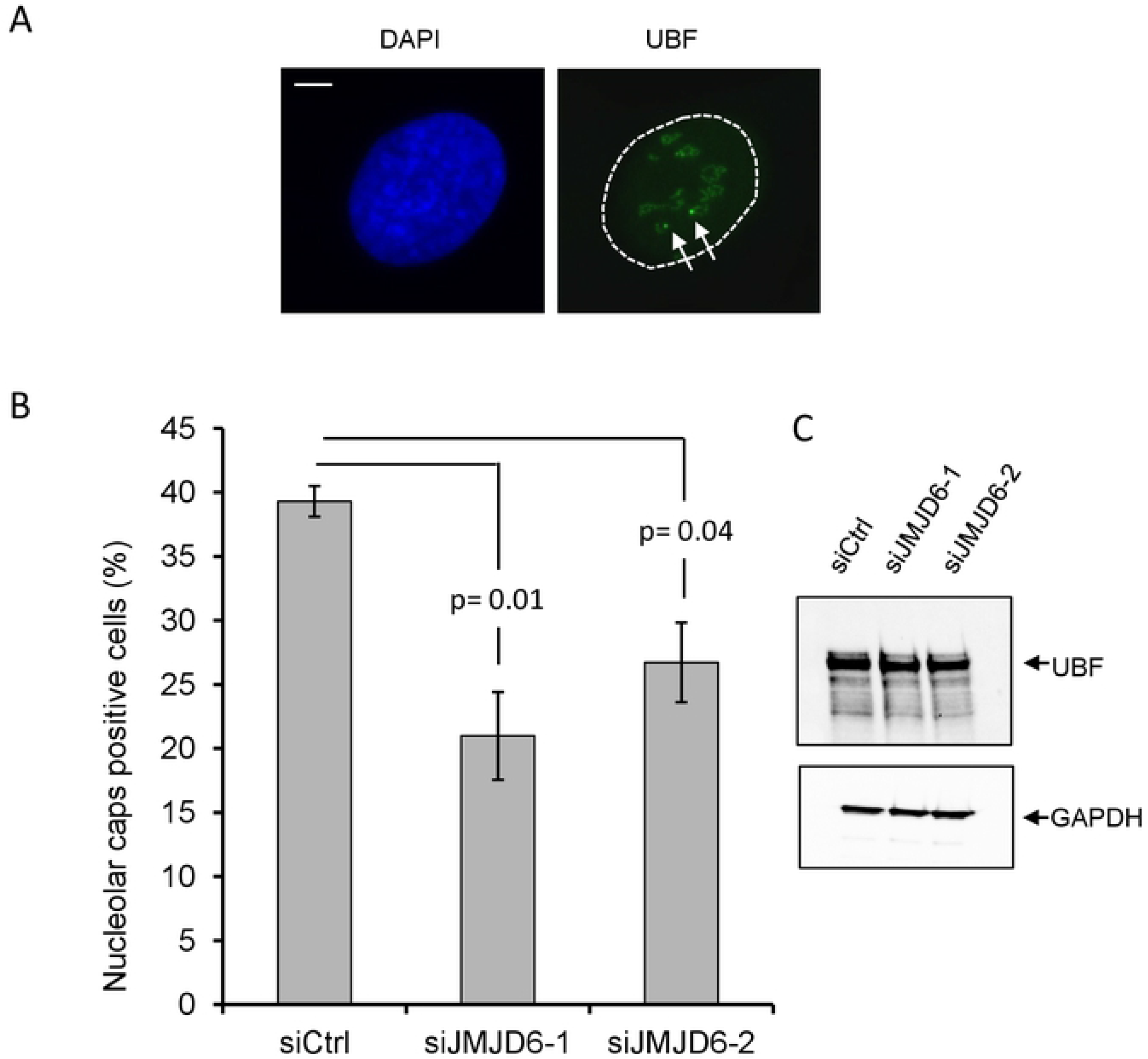
Nucleolar caps formation in response to DNA damage. A. Images from OHTam-treated DIVA cells showing nucleolar caps revealed by UBF foci formation at the nucleolus periphery. The arrows point towards nucleolar caps. Scale bar 5 µm. B. Quantification of nucleolar caps positive cells after JMJD6 depletion. The mean and standard deviation from three independent experiments are shown. For each conditions a minimum of 100 cells were counted. C. Western blot analysis of UBF expression after siRNA-induced JMJD6 depletion.

Previous studies showed the importance of ATM signaling for the generation of nucleolar caps and transcription inhibition [9,13]. We thus investigated the relationship between JMJD6 and ATM in the pathway leading to nucleolar caps. First, we tested the activation of ATM in the different cell lines in response to IR. As shown in the Figure 8A, ATM was normally activated in WT- and JMJD6-KO cell lines in response to IR. Then, we tested whether ATM signaling and JMJD6 were in the same pathway for the generation of nucleolar caps. As previously described for JMJD6 knockdown cells (Figure 7B), JMJD6 KO cells were defective for nucleolar caps formation. JMJD6-KO cells complemented with the WT form of JMJD6 presented more nucleolar caps, this increase being sensitive to ATM inhibition (Figure 8B). Thus, these data indicate that JMJD6 and ATM are in the same pathway. Considering that ATM activation is normal upon JMJD6 inactivation and that the ATM-dependent transcription inhibition process is not abolished upon JMJD6 knock-down, these data suggest that JMJD6 lies downstream of ATM in the pathway leading to nucleolar caps formation

**Figure 8.**
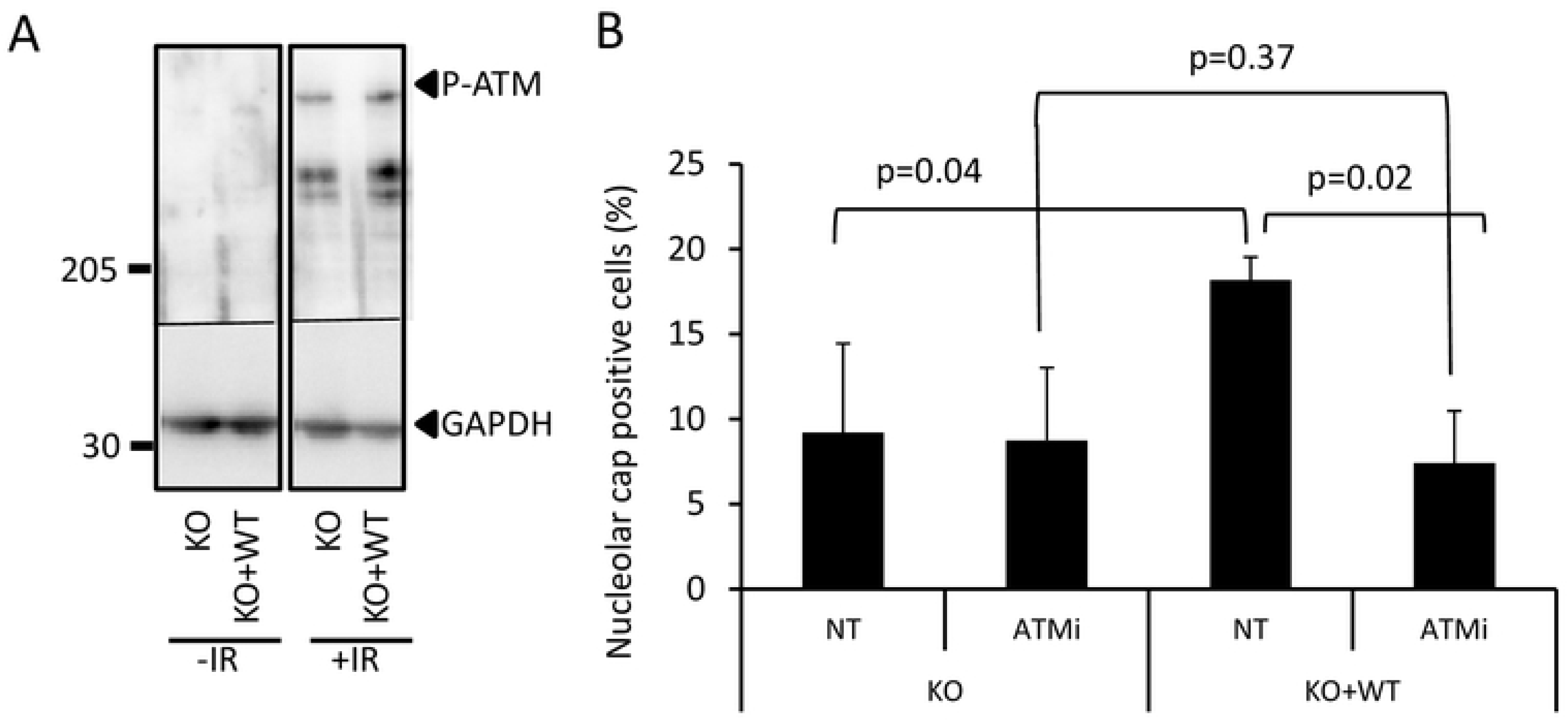
ATM activation in JMJD6 defective cells. A. Western blot analysis of ATM activation in response to IR (20 Gy 1h post IR) performed by the detection of the phosphorylated form of ATM. B. Dependence on ATM activity for the formation of nucleolar caps in response to IR. Cells were treated or not with the ATM inhibitor. A minimum of 100 cells were counted for each conditions in each independent experiments. Results are the mean +/− standard deviation from three independent experiments. The p values of the difference between the indicated samples are indicated (Student t test).

### JMJD6 affects the recruitment of NBS1 into the nucleolus in response to DNA damage

DNA damage in the nucleolus need to be repaired before restart of the transcription. To repair DSB cells can use HR or NHEJ. The most rapid pathway to repair DSB is the use of NHEJ but for complex lesions or DSB requiring faithful repair cells can use homologous recombination. Both process are influenced by NBS1 via its involvement in the MRN complex. Because we observed that JMJD6 interact with TCOF we wondered whether JMJD6 could alter NBS1 recruitment into the nucleolus.

To test whether JMJD6 exerts a similar role to TCOF in the control of NBS1 localisation, we assayed the presence of NBS1-GFP foci in the nucleolus of JMJD6 KO cells in response to IR (Figure 9A). As expected, we observed an increase in NBS1 foci in the nucleolus of control cells in response to IR exposure. In contrast, IR-induced nucleolar localization of NBS1 was strongly decreased in JMJD6 KO cells, an effect which was reversed when these cells were complemented with JMJD6 but not in the cells complemented with the inactive form of JMJD6 (Figure 9B). These results show that JMJD6 and its enzymatic activity are required for the recruitment of NBS1 to the nucleolus in response to DNA damage, providing a direct link between JMJD6 and the presence of DNA repair factors in the nucleolus.

**Figure 9.**
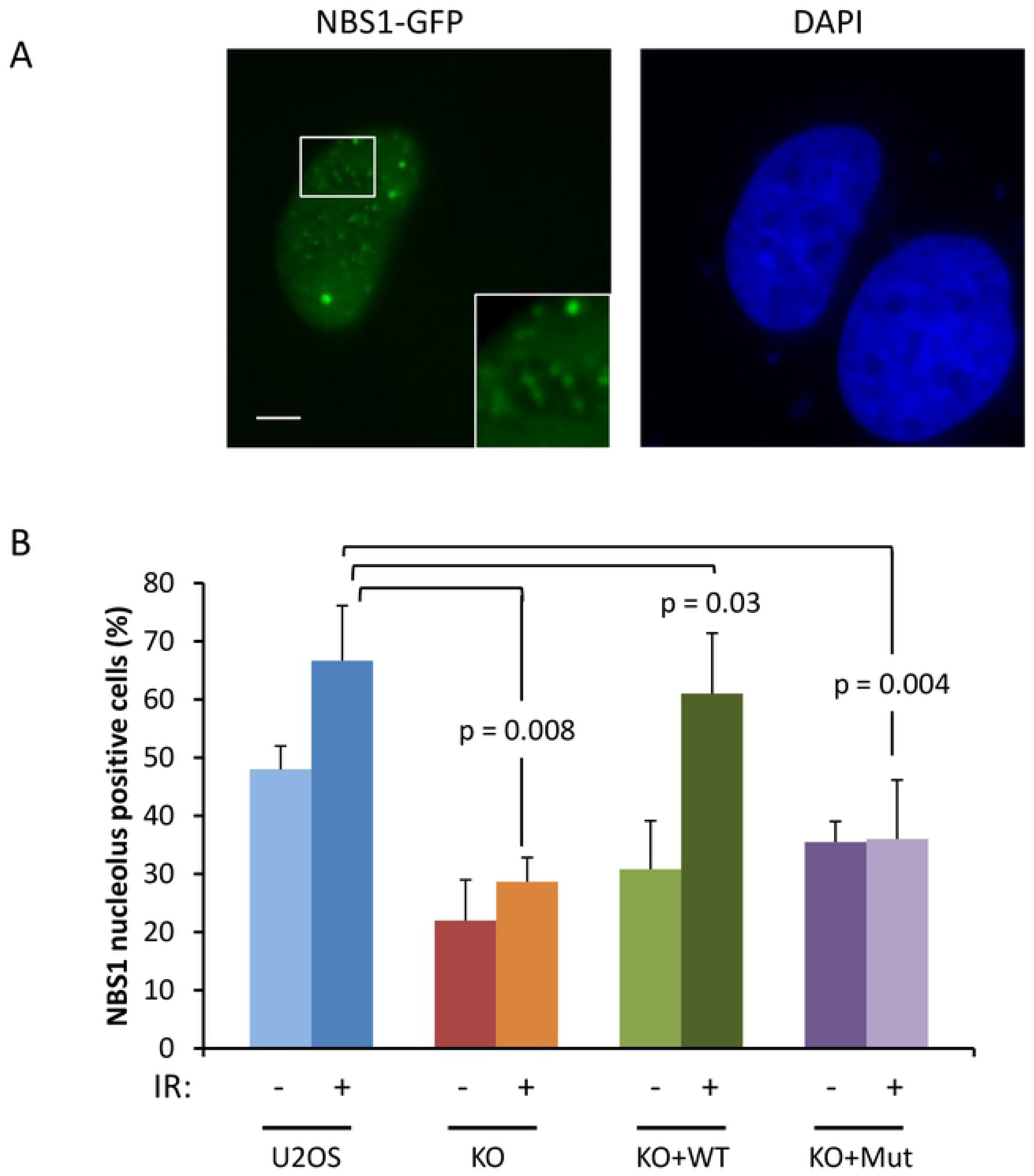
NBS1 localization in nucleoli after DNA damage is dependent on JMJD6. A. Representative images of U2OS cells expressing NBS1-GFP after IR exposure (5 Gy). Insert showing the presence of NBS1 foci in nucleolus. Scale bar 5 µm. B. Quantification of the percentage of U2OS, JMJD6-KO and JMJD6-KO-complemented with wild type JMJD6 (KO+WT) and JMJD6-KO-complemented with an inactive form of JMJD6 (KO+Mut) cell lines expressing NBS1-GFP that exhibit NBS1 nucleolar foci without and after IR exposure. Results are the mean +/− standard deviation from three independent experiments. In each independent experiment more than 100 transfected cells were counted for each point. The p values of the difference to the U2OS cell line after IR are indicated (Student t test).

## Discussion

Here, we demonstrate for the first time that the JMJD6 histone demethylase is important for the response to DSBs occurring in rDNA repeats in the nucleolus. This relies on four independent lines of evidence: first, JMJD6 is mobilized upon DNA damages induced in the nucleolus. Second, JMJD6 depleted cells are more sensitive to DSBs specifically occurring in rDNA. Third, JMJD6 depleted cells harbour genetic instability at rDNA loci upon DNA break induction. Fourth, JMD6-depleted cells are defective in the formation of nucleolar caps in which repair of persistent DSBs by homologous recombination is generally thought to occur.

Our data suggests that JMJD6 function is intimately linked to the role of Treacle. Indeed, we found a physical interaction between JMJD6 and Treacle. Moreover, ChIP experiments indicated that JMJD6 is present at the rDNA before break induction, as is Treacle[26]. Finally, again like Treacle, we found that JMJD6 is important for the full recruitment of NBS1 in the nucleolus. Interestingly, in the absence of JMJD6, transcription repression at rDNA still occurs, indicating that there is enough NBS1 to mediate transcriptional repression. However, there are defects of NBS1-dependent processes, such as formation of nucleolar caps. Our data thus suggest that JMJD6 is specifically required for NBS1 to participate in DNA repair processes occurring at the rDNA. NBS1 within the MRN complex is involved in the two main repair processes of DSB repair, NHEJ and HR [29]. The MRN complex mediates DNA resection at DSBs, a process which is required for nucleolar caps formation [28]. This probably explains why nucleolar caps formation is defective upon JMJD6 inhibition. However, it is also possible that NHEJ is also defective upon JMJD6 depletion. Indeed, we found that JMJD6 depletion affects NHEJ in the nucleoplasm, as measured using reporter substrate stably integrated in DNA but outside the rDNA (data not shown). Defects into the two main mechanisms of DSB repair in the nucleolus is probably responsible for the genetic instability of rDNA arrays we observed in JMJD6 depleted cells, with a decreased number of rDNA arrays and complex rearrangements of repeats (Figure 3).

The fact that actinomycin D treatment induced nucleolar caps led to the proposal that their formation was a direct consequence of transcriptional inhibition [30]. However, we show here an example of a protein whose depletion leads to the uncoupling between transcription inhibition and nucleolar caps formation, indicating that cap formation is an active process. Strikingly, a similar conclusion was drawn from a recent study which shows that the LINC complex is important for nucleolar caps formation downstream of transcriptional inhibition [28]. These observations indicate that the formation of nucleolar caps is not a consequence of transcription inhibition *per se* but requires specific factors.

We also found that rDNA transcription is lower upon DNA break induction in JMJD6 depleted cells compared to control cells. This could be due to a direct role of JMJD6 in allowing rDNA transcription recovery following DSB repair by controlling the interaction between Treacle and NBS1. Alternatively, it may be the consequence of a role of JMJD6 in NHEJ-mediated repair of rDNA breaks, since rDNA transcription silencing in response to DNA damage is exacerbated upon inhibition of NHEJ [9]. In addition, our mass spectrometry data reveal JMJD6-PHRF1 interactions in etoposide treated cells (see the complete list of JMJD6 interactors with the link provided in Materials and Methods section). PHRF1 has been described to influence DSB repair by NHEJ and to interact with NBS1 [31]. How this would be related to the function of JMJD6 in nucleolar caps formation is unclear and could reflect independent roles of JMJD6 in the management of DNA breaks occurring in rDNA.

Although we studied JMJD6 involvement in the repair of DSBs occurring in rDNA, we cannot rule out that JMJD6 is also involved in the repair of damage occurring at other genomic locations. Strikingly, such an effect was recently described by Huo et al. [21] However, we obtained results contradictory to them in particular concerning the sensitivity to IR in JMJD6 defective cells. This is probably due to the fact that we examined IR sensitivity in KO cells whereas they used KD performed after siRNA. The study by Huo et al showed recruitment of GFP-JMJD6 in the nucleus after laser induced DNA damage but also in the nucleolus but the reason of such observation and its consequences were not addressed. In addition, they showed that the phenotype observed were independent of JMJD6 activity whereas in our study they are showing a direct link between JMJD6 and its involvement in DNA damage response at ribosomal DNA. Finally, they identified SIRT1 as interacting with JMJD6 that we do not observed in our mass spectrometry data. Several reasons could explain such discrepancy such as the use of overexpressed N-terminus Flag-tagged JMJD6 in Huo et al. study whereas we use endogenously C-terminus Flag-Strep tagged JMJD6 [21]. Whether JMJD6 plays a general role in DNA damage response and/or repair, or whether it is important for repair of specific loci clearly merits further investigations. Potential candidate loci could be DNA elements with features similar to rDNA arrays, with a repetitive nature and requiring sequence conservation. In summary, we identified here a new factor participating in the maintenance of rDNA integrity. The importance of such mechanism is highlighted by the increasing number of studies showing the involvement of rDNA repeats integrity in a number of disease such as cancer, neurological- and aging-associated diseases most of them in relationship with gene products associated with the genome maintenance [4,8].

## Materials and Methods

### Cell culture conditions

U2OS cell line, and subclones were cultured in DMEM medium supplemented with 10% foetal calf serum, sodium pyruvate and antibiotics (penicillin/streptomycin) in a humidified atmosphere with 5% CO2. K562 cell line was cultured in RPMI 1640 medium supplemented with foetal calf serum and antibiotics (penicillin/streptomycin).

### Generation of JMJD6 deficient and tagged cell lines

U2OS cells were made defective for JMJD6 by using CRISPR technology as described in [22]. Homozygote knocked out cell lines were checked by sequencing and Western blot for JMJD6 expression. A KO cell line was transfected with a plasmid coding for a V5-tagged form of JMJD6 (KO+WT) or a JMJD6 catalytically inactive form (H187A-D189A-H273A) (KO+Mut) (gift of Dr M Le Romancer, Lyon, France) and stable clones selected in order to complement the JMJD6 deficiency. A Flag-Strep tag was inserted in K562 cells at the C-terminus of JMJD6 using CRISPR technology as described [22,32].

### High throughput microscopy

U2OS cells (7500) were transfected with 10nM siRNA using INTERFERIN (Ozyme) according to the manufacturer’s instructions and seeded in 96 well plates (Perkin Elmer). Two days after transfection cells were exposed to ionizing radiation (8 Gy) then fixed with formaldehyde (4% in PBS). Cells were permeabilized with triton X100 (0.5 % in PBS) for 5 min, stained with stained with γH2AX antibody (1/500) diluted in PBS-BSA 1% and Alexa Fluor 488 anti mouse (Thermofisher). Acquisition was performed on at least 1000 cells per well (3 wells per condition) with 20X objective with Harmony Imaging Software 4.1 (Perkin Elmer). Image analysis was pursued using Colombus 2.5.0 software (Perkin Elmer) to quantify γH2AX spot intensity.

### 5FUrd incorporation measurement

Transcription of rDNA was monitored by revealing the incorporation of 5-Fluoro-Uridine in Nascent RNA (5-FUrd, Sigma). Cells were incubated in presence of 2mM 5FUrd for 20 minutes followed by fixation in 4% FA (Sigma). They were permeabilized with triton X100 (0.5 % in PBS) for 5 min, then stained using an anti BrdU antibody (Sigma, B2531) and Alexa Fluor 488 anti mouse (Thermofisher) diluted 1/500 in PBS-BSA 1% and 0.4 U.mL^-1^ RNAsin (Promega). We visualized and quantified 5-FUrd incorporation by using high content imaging device (OPERETTA, Perkin Elmer). All imunofluorescence steps were performed at 4°C. Image analysis was pursued using colombus software to measure 5-FUrd spot intensity.

### Laser-induced DNA damage on living cells

The system used to perform laser-induced DNA damage has been previously described in details in [33]. Briefly, the system is composed of an inverted microscope (DMI6000B; Leica) equipped with a temperature controller and a CO2 flow system. DNA damage was generated on nucleus with a green pulsed laser (532 nm). The beam was focused with a 100x NA 1.4 immersion objective (Leica). Images were acquired with a cooled charge-coupled device camera (CoolSNAP HQ2). The system was driven by Metamorph software. U2OS cells were transfected with plasmid expressing JMJD6-GFP in a 2-well chamber (Labtek) in 1 ml of OptiMEM medium without red phenol. Images were recorded using the Metamorph software package (MDS Analytical Technologies).

### Detection of nucleolar caps

DIvA cells were seeded on glass coverslips, transfected with the different siRNA (10nM final concentration) according to the manufacturer’s instructions with Jet Pei (Ozyme, France). Two days later, cells were treated with hydroxy-tamoxifen during 4h to generate DNA DSB [27]. Cells were fixed with formaldehyde (3.7% in PBS) permeabilized with triton X100 (0.5 % in PBS) for 5 min, washed, then non-specific binding saturated with PBS-BSA (3%) for 1h and incubated with anti UBF antibody (1/500) in PBS-BSA 0.5% for 1h then with secondary antibody coupled to fluorophore (alexa 488) and stained with DAPI. In the case of ionizing radiation exposure, cells were treated with 20 Gy and fixed 6h post IR exposure. When necessary cells were treated with the ATM inhibitor (KU55933) (10 µM final concentration) 1h prior to IR exposure.

### Proximity Ligation Assay (PLA)

The cells were seeded onto glass coverslips, treated (5 Gy) fixed (1h post IR) with formaldehyde (3.7% in PBS) and the PLA assays (Duolink PLA technology, Sigma-Aldrich, DUO92014) performed using manufacturer’s protocol with antibody staining performed as in the standard immunofluorescence procedure with antibodies against V5 tag and Treacle or NBS1 and Treacle

### Imaging

Images were collected with a microscope (DM5000; Leica) equipped with a charge-coupled device camera (CoolSNAP ES; Roper Scientific) and a 100x objective (HCX PI APO ON:1.4-0.7) and SEMROCK filters. Acquisition software and image processing used the MetaMorph software package (Molecular Devices). Confocal imaging was performed on LSM880 microscope (Zeiss), Z-stacks of fluorescent images were captured and analyzed using ZEN software with a 63x immersion oil objective (Plan-Apochromat ON:1.4).

### Immublotting detection

Whole cell extracts were prepared in Laemmli buffer (Tris HCl 62.5mM, SDS1%, Mercaptoethanol 5%, glycerol 25% and bromophenol blue). Samples were sonicated and heated at 95°C for 5 minutes before their separation on a 4-15% gradient SDS PAGE gel (Biorad, 4-15%). Proteins were transferred from gel to nitrocellulose membrane using semi-dry method. Once blocked in PBS-Tween (5%) plus 10% skimmed milk, membrane were firstly immunoblotted with primary antibody diluted in 2% milk PBS-T (generally 1/1000 except for anti H3 whose dilution used is 1/10000) at 4°C overnight and finally immunobloted with secondary antibody coupled to HRP (Sigma) diluted in 2% milk PBS-T at room-temperature for 1 hour. After several washes in PBS-T, proteins were detected using ECL lumi-light plus (Roche) and images acquired using camera system (Chemitouch Biorad).

### Clonogenic assay

Cells were seeded at 500 cells per well in 6 well plates and let to attach overnight. Plates were irradiated using a Biobeam 8000 (^137^Cs source) (Anexplo service, Rangueil, Toulouse, France). Plates were kept in cell incubator for 10 days then stained with crystal violet. Colonies of more than 50 cells were counted.

For purpose of testing cell sensitivity to DSB in rDNA, U2OS cells (1 x 10^6^) were transfected with siRNA (1µM) by electroporation (4D-Nucleofactor, Amaxa) and seeded at 1.10^5^ cells per well in 6-well plates. 48h after transfection, cells were transfected by JetPEI (Ozyme) according to manufacturer’s instructions with either an empty px330 plasmid (Addgene) coding only for Cas9 or a px330 plasmid coding for Cas9 and a sgRNA targeting rDNA [13], together with px330 coding for a sgRNA targeting ATP1A1 gene together with a donor plasmid to generate a mutated ATP1A1 gene coding for an ouabain insensitive Na^+^/K^+^ ATPase [22]. 48 hours after plasmids transfection, ouabain (0.7 µM, Sigma, France) was added to the culture medium to select transfected cells in which CRISPR-induced cleavage and mutation insertion happened. Two weeks later cells were fixed and stained with crystal violet solution and the colonies counted.

### Purification and mass spectrometry analysis of endogenous JMJD6 interactome

K562 cells expressing 3xFlag-2xStrep tag at the C-terminus of endogenous JMJD6 were amplified (1.5 x10^9^ cells), nuclear cell extracts prepared and used to perform tandem affinity purification as described in [34].

Briefly, nuclear extracts [35] were adjusted to 0.1% Tween-20, and ultracentrifuged at 100,000 g for 45 min. Extracts were precleared, then 250 ul anti-FLAG M2 affinity resin (Sigma) was added for 2 hr at 4°C. The beads were then washed in Poly-Prep columns (Bio-Rad) buffer #1 (20 mM HEPES-KOH[pH 7.9], 10% glycerol, 300 mM KCl, 0.1% Tween 20, 1 mM DTT, Halt protease and phosphatase inhibitor cocktail [Pierce]) followed buffer #2 (20 mM HEPES-KOH [pH 7.9], 10% glycerol, 150 mM KCl, 0.1% Tween 20, 1mMDTT, Halt protease and phosphatase inhibitor cocktail [Pierce]). Complexes were eluted with buffer #2 supplemented with 150 ug/ml 3xFLAG peptide (Sigma) for 1 hr at 4°C. Next, fractions were mixed with 125 ul Strep-Tactin Sepharose (IBA) affinity matrix for 1 hr at 4°C, and the beads were washed with buffer #2 in Poly-Prep columns (Bio-Rad). Complexes were eluted in two fractions with buffer #2 supplemented with 2.5mM D-biotin, flash frozen in liquid nitrogen, and stored at −80°C. Typically, 15 ul of the first elution (3% of total) was loaded on NuPAGE 4%–12% Bis-Tris gels (Life Technologies) and analyzed by silver staining.

For mass spectrometry analysis, fractions were loaded on gel and migrated for about 1 cm, then stained with sypro ruby red and a gel slice containing the entire fraction was cut for in-gel trypsin digestion and analysis on a LC-MS/MS apparatus (Thermo scientific Orbitrap Fusion) at the Quebec Genome Center.

### MS data archiving

All MS files used in this study were deposited at MassIVE (http://massive.ucsd.edu) and at ProteomeXchange (http://www.proteomexchange.org/). They were assigned the identifiers MassIVE MSV000083409 and PXD012603. The password to access these files until publication is “JMJD6”.

### Measurement of genetic instability at rDNA

Cells were seeded on glass coverslips and irradiated at 2 Gy and let recover for 24h before adding colcemid (0.1 µg/ml) for 3h to enrich the cell population in metaphases. Then cells were fixed with formaldehyde (3.7% in PBS) permeabilized with Triton X100 (0.5 % in PBS) for 5 min, washed, then non-specific binding saturated with PBS-BSA (3%) for 1h and incubated with anti UBF antibody (1/500) in PBS-BSA 0.5% for 1h then with secondary antibody coupled to fluorophore (alexa 488) and stained with DAPI.

### DNA FISH combing on ribosomal DNA

Cells were transfected with control siRNA (siCtrl), or two different siRNAs directed against JMJD6 (siJMJD6-1 and siJMJD6-2) and 48h later induced or not for DSB by treatment with OHTam during 24h. At the end of the treatment cells were harvested and included in low melting agarose and sent to the Genomic Vision Company for combing and FISH. DNA fibers stained as represented in figure 9 were then detected using web interface provided by Genomic Vision company. Identification and counting of rearrangement events were estimated using a script designed to count the total number of rDNA units and assessing the the occurence of changes when comparing one unit to the next. GitHub https://github.com/LegubeDNAREPAIR/rDNA/blob/master/get_break_event.py

### Chromatin Immunoprecipitation (ChIP)

Cells were double crosslinked with DMA (0.25%, 45 min), washed with PBS then with formaldehyde (1%, 20min) and ChIP was performed as described (Mattera courilleau 2010) using 200 µg of chromatin. Briefly, nuclei were prepared and sonicated to obtain DNA fragments of about 500–1000 bp. Following preclearing and blocking steps, samples were incubated overnight at 4°C with specific antibodies (anti V5) or without antibody (mock) as negative control. Immune complexes were then recovered by incubating the samples with blocked protein A/protein G beads for 2 h at 4°C on a rotating wheel. After extensive washing, crosslink was reversed by adding RNase A to the samples and incubating overnight at 65°C. After a 1h30 proteinase K treatment, DNA was purified with the GFX PCR kit (Amersham), and analysed by Q-PCR on a CFX96 Real-time system device (BioRad) using the IQ SYBR Supermix (BioRad Laboratories, Marnes-la-Coquette, France), according to the manufacturer’s instructions. All samples were analyzed in triplicates.

### List of antibodies used

Anti JMJD6 (Santa Cruz, sc28349), anti UBF (Bethyl, A301-859A), anti histone H3 (Abcam, Ab1791), anti BrdU (Sigma, B2531), anti gamma H2AX (Cell signaling, 9718(20E3)), anti GAPDH (Chemicon, MAb374), anti myc (Santa Cruz, sc-40), anti Treacle (Santa Cruz, sc374536), anti V5 (Cell Signaling, 12032), anti NBS1 (Sigma, PLA0179), anti phospho ATM (S1981) (Cell signaling, 10H11.E12).

### List of oligonucleotides used

siRNA control: CTTACGCTGAGTACTTCGA; siRNA JMJD6-1: CAGCUAUGGUGAACACCCUAA; siRNA JMJD6-2: CCAAAGUUAUCAAGGAAAU sgRNA:rDNAtarget: GCCTTCTCTAGCGATCTGAG sgRNA: JMJD6KO: GAGCAAGAAGCGCATCCGCG sgRNA: JMJD6 tag: CCAGGTGACCCAGCAAGGCT

### Statistical analysis

Data obtained with operetta device were analysed using Wilcoxon Mann-Whitney test. For data with lower number of values a Student t test was performed (clonogenic assay).

## Acknowledgements

We thank Thomas Mangeat for his assistance with the laser track experiments and Magali Suzanne for assistance with the confocal experiments. We are grateful to the Non-Invasive Exploration service-US006/CREFRE INSERM/UPS for giving us the access to the irradiator Biobeam 8000 (Rangueil, Toulouse). We acknowledge the Toulouse Regional Imaging - Light Imaging Toulouse CBI Platform for technical assistance in high content microscopy. We thank Florence Larminat for her critical reading of the manuscript.

## Supporting information legends

**Figure supplement S1. Laser induced DNA damage on living cells.**

A. U2OS cells expressing 53BP1-GFP. Dotted red line indicated laser irradiation. The time post irradiation is indicated above images. B same as in A with cells expressing GFP protein. C. additional cells expressing JMJD6-GFP as in Figure 1.

**Figure supplement S2. JMJD6 recruitment at DSB assessed by ChIP**

Chromatin immunoprecipitation (ChIP) in DiVA cell line transfected with tagged JMJD6-V5. ChIP results are expressed as fold enrichment compared with signal obtained on beta-actin set at 1. Mock: no antibody; V5: anti V5 antibody. Results are expressed as mean+/− sd of triplicate qPCR.

**Figure supplement S3: JMJD6 KO cell lines present increased sensitivity to ionizing radiations.**

A. U2OS cells or two clones of U2OS cells inactivated for JMJD6 expression (KO and KO#2) were exposed to ionizing radiations and subjected to a clonogenic assay. The mean and standard deviation from three independent experiments are shown. B. Western blot analysis for JMJD6 expression using JMJD6 and Histone H3 antibodies. The bar indicates that the original image was cut. The stars indicate two non specific bands detected by the anti-JMJD6 antibody. C. same as in A with JMJD6-KO cell line complemented with JMJD6. D same as in B.

**Figure supplement S4. Recruitment of JMJD6 mutant at DNA damages.**

U2OS cells expressing the inactive JMJD6-Mut-GFP. Dotted red line indicated laser irradiation. The time post irradiation is indicated above images.

**Figure supplement S5. Analysis of rDNA rearrangements in JMJD6-depleted cells in absence and after DSB induction.**

Results of the second DNA FISH experiment performed totally independently from the first experiment presented in figure 9. Results are NT: not treated, OHT: induction of DSB after OHTam treatment as reported in materials and methods.

**Figure supplement S6. JMJD6 colocalizes with Treacle in nucleolus.**

Image of U2OS cells expressing V5-tagged-JMJD6 exposed to ionizing radiations at 5 Gy (1 h post-IR) analysed by confocal microscopy using anti-V5 and anti-Treacle antibodies. Confocal line profile showing in red Treacle, in cyan JMJD6-V5, and DAPI in white. Note the correspondence of the peaks for Treacle and JMJD6.

**Figure supplement S7. JMJD6 influences the NBS1-Treacle interaction.**

Independent experiment from Figure 5 monitoring the NBS1-Treacle interaction after DNA damage.

**Figure supplement S8. JMJD6 colocalizes with Treacle and NBS1 in nucleolus.**

Images of U2OS cells expressing V5-tagged-JMJD6 transfected with NBS1-GFP and exposed to ionizing radiations at 5 Gy (1 h post-IR) analysed by confocal microscopy using anti-V5 or anti-Treacle antibodies. Confocal line profile showing in red Treacle, in cyan JMJD6-V5, in green NBS1-GFP and DAPI in white. Note the correspondence of the peaks for Treacle and JMJD6 and NBS1.

**Figure supplement S9: cell cycle repartition of JMJD6 depleted cells**

U2OS cells were transfected with the indicated siRNA. 48 hours later, they were treated with EdU for 30 minutes, then harvested and analysed by flow cytometry for propidium iodide staining and EdU incorporation. The repartition of transfected cells in the various phases of the cell cycle is indicated

